# Herbarium-based measurements are reliable predictors of fresh plant traits in Neotropical Myrtaceae

**DOI:** 10.64898/2026.03.26.714626

**Authors:** Yacov Kilsztajn, Lázaro Henrique Soares de Moraes Conceição, Thais Vasconcelos, Carolyn Elinore Barnes Proença, Vanessa Graziele Staggemeier

## Abstract

**Premise:** Herbarium specimens are increasingly used to extract morphological traits for ecological and evolutionary studies, yet the effects of tissue desiccation on trait measurements remain poorly understood. Here, we tested whether higher tissue water content leads to greater measurement changes after herborization (H1) and whether fresh trait values can be reliably predicted from herbarium measurements (H2).

**Methods:** We evaluated the reliability of herbarium-based measurements by comparing fresh and dried traits of leaves, flowers, fleshy fruits, and seeds across 262 individuals representing 133 Neotropical Myrtaceae species. Phylogenetic least square models and machine-learning regressions were used to test H1 and H2.

**Results:** Leaves and flowers generally shrank after herborization, fruits size metrics tended to increase, and seeds were largely unaffected. Water content was significantly associated with the magnitude of herborization effects in flowers and some leaf and seed traits. Fresh trait values were accurately predicted from herbarium measurements. Prediction errors were lowest for leaf traits, followed by fruits, flowers, and seeds.

**Discussion:** These results partially support H1 and support H2, indicating that herbarium specimens can be reliably used for trait analyses when organ-specific responses are considered, providing a practical framework to account for potential desiccation bias in functional trait research.

## INTRODUCTION

Herbarium specimens have long served as an important resource in plant taxonomy, as botanists rely on dried plant specimens to describe new species and establish taxonomic classifications (Morton, 1981). These preserved materials also are an invaluable historical record of plant diversity and geographical distribution (Mandrioli, 2023; Marín-Rodulfo et al., 2024). In recent decades, the utility of herbarium data has expanded well beyond taxonomy. Morphological measurements from preserved specimens have increasingly been used to address ecological and evolutionary questions (Jaroszynska et al., 2023; Velásquez-Puentes et al., 2023; Wilde et al., 2023). This broader application has coincided with the digitalization of herbarium collections and the growing accessibility of specimen metadata and high-resolution images (Soltis, 2017).

As a consequence, the combination of herbarium specimens with modern analytical tools is not just improving our understanding of plant biology, but also creating a new paradigm for biodiversity research, creating a wealth of accessible historical data. For instance, recent advances in artificial intelligence and computer vision have enabled the automated extraction of morphological traits from digitized herbarium specimens (e.g., Weaver et al., 2020; Hussein et al., 2021; Guo et al., 2024; Vasconcelos et al., 2025). These innovations have fueled large-scale studies that integrate thousands of specimens across wide spatial and temporal scales, offering unprecedented opportunities to explore functional trait variation in an ecological and evolutionary context.

However, the use of herbarium specimens in such analyses introduces new challenges related to data accuracy and potential measurement biases. One such bias arises from the desiccation during specimen preparation. As plant tissues dry they usually shrink, potentially altering the size and shape of morphological traits. Although this phenomenon has been acknowledged, few studies have systematically assessed its impact on trait measurements in herbarium specimens, with most of them focusing primarily on vegetative characters, such as leaf traits (e.g., Queenborough and Porras, 2014; Perez et al., 2020; Blonder et al., 2012). The extent of shrinkage and deformation is likely to vary among plant organs and may depend on their initial water content (Tomaszewski and Górzkowska, 2016). Structures with higher water content, such as flowers and fleshy fruits (Roddy et al., 2023), are expected to shrink more during drying than denser organs, like leaves and seeds.

No study to date has explicitly tested the influence of herborization on the accuracy of trait measurements across a range of plant structures. We explore this topic by using the Neotropical clade (*sensu* Vasconcelos et al., 2017) of the tribe Myrteae (Myrtaceae) as a model group (hereafter Neotropical Myrtaceae), which includes around 50 genera and 2,700 species (NMWG et al., 2024). This group is highly variable in terms of vegetative and reproductive traits, including membranaceous to leathery leaves of different sizes, very small to large flowers arranged either solitarily or in inflorescences bearing few to dozens of flowers, and a wide variety of fruit and seed morphologies (POWO, 2024). Such morphological diversity makes Neotropical Myrtaceae an ideal system to investigate the effects of herborization on trait measurements, while also enabling broader inferences applicable to other plant clades, particularly fleshy-fruited woody taxa.

We quantify the divergence between trait measurements obtained from herbarium specimens and those measured in fresh material for Neotropical Myrtaceae species. We explicitly test two hypotheses: (H1) the magnitude of size change caused by herborization increases with the water content of the plant organ; and (H2) fresh trait values can be reliably predicted from herbarium-based measurements.

## METHODS

### Field and herbarium sampling

Samplings were conducted in 26 localities across Brazil, including areas belonging to three different biomes (Olson et al., 2001): Tropical and subtropical grasslands, savannas, and shrublands (n=6), Tropical and subtropical moist broadleaf forests (n=14), and desert and xeric shrublands (n=6) (Fig. S1). We targeted reproductive individuals of Neotropical Myrtaceae species bearing either flowers and/or mature fruits. Whenever possible, three to five individuals per species were sampled, and two structures per organ were measured for each specimen. In total, we sampled 262 individuals, from 133 species across 14 genera and 7 subtribes (Table S1).

We quantified a set of morphological traits, including leaf area, leaf length/width ratio, leaf circularity index, leaf mass per area, petal length, flower length and width (petal-to-petal and stamen-to-stamen), hypanthium length and width, and fruit and seed size length and width (measurements followed Pérez-Harguindeguy et al., 2016; see Fig. S2 for details on flower measurements). All traits were measured from photographs of fresh structures flattened in an acrylic press with a 5 cm scale positioned alongside the specimens, using ImageJ (Schneider et al., 2012). Fruit length and width were also measured prior to pressing (except for 7 specimens for logistic reasons), as flattening alters the dimensions of mature fleshy fruits. Measurements from pressed fruits were therefore used only to disentangle the effects of pressing and drying on fruit morphology.

After fresh measurements, all samples were subjected to standard herborization procedures. Plant material was carefully arranged between sheets of newspaper and cardboard to ensure uniform flattening and moisture absorption. The layered material was placed in a plant press and evenly compressed using adjustable ropes, with pressure manually adjusted according to organ thickness and tissue consistency to minimize deformation. Pressed specimens were dried in a closed drying chamber for four to seven days. After drying, all morphological measurements were repeated using the same photographic and analytical protocols applied to fresh material. Voucher specimens were prepared in parallel and deposited in the following herbaria: ALCB, CEPEC, IAN, MBM, MICH, MG, PMSP, RB, SPF, UB, and UFRN (acronyms according to Thiers, continuously updated). Fresh and dry mass were recorded for each plant structure (leaves, flowers, fruits, and seeds; petioles and peduncles were retained for mass measurements) using a portable balance (Brifit Mini Scale, 0.01 g precision), allowing estimation of water content. Leaf mass per area was calculated using a constant dry mass value for both fresh and dry leaves, divided by leaf area measured in the fresh and dry states, respectively, following Pérez-Harguindeguy et al. (2016).

### Herborization effect and water content

To quantify the morphological shifts induced by drying, we calculated indices based on paired measurements of fresh and dried materials. First, we estimated the water content (*WC*) of each plant structure (leaves, flowers, fruits, and seeds) for every sampled specimen as the proportional difference between fresh (*M*_*fresh*_) and dried mass (*M*_*dried*_).

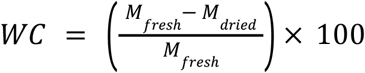

Second, we quantified the herborization effect (*I*_*h*_) for each trait as the ratio between the value measured in dried material (*T*_*dried*_) and its fresh counterpart (*T*_*fresh*_), minus one.

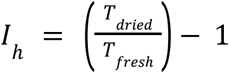

This index represents the proportional change in trait dimensions after herborization, where positive values indicate expansion and negative values indicate shrinkage.

For fruits (that are three-dimensional organs), we applied a two-step process in order to distinguish between the effect of pressing (mechanical compression) and drying (desiccation). Specifically, the pressing effect was calculated as the ratio between the measurements obtained from newly collected, fresh fruits and the measurements of these fruits pressed (prior to drying), minus one. The drying effect, in turn, was calculated as the ratio between the measurements obtained from the fresh pressed fruits and the desiccated pressed fruits, minus one. By comparing these indices, we were able to partition the total morphological distortion into its mechanical (pressing) and desiccation components. In both cases, positive values indicate expansion, whereas negative values indicate shrinkage.

We tested whether tissue water content explains variation in herborization effect using phylogenetic generalized least squares (PGLS) models fitted to species mean trait values. Analyses were based on the most comprehensive dated phylogeny currently available for Neotropical Myrtaceae (NMWG et al., 2024). Models were fitted separately for leaves, flowers, fruits, and seeds using a maximum-likelihood PGLS framework with Pagel’s λ estimated from the data, implemented in the R package phylolm (Ho et al., 2016).

To incorporate species lacking phylogenetic placement in the backbone phylogeny (41 species out of 133; Table S1), we generated a set of alternative trees by randomly inserting missing taxa within their most likely clades using the R package ape (Paradis and Schliep, 2019). Insertions followed a hierarchical taxonomic procedure: when information on genus sections was available (Table S1), species were randomly inserted along branches within the clade defined by congeners belonging to the same section; otherwise, insertions were performed within the genus. This procedure was repeated to generate 100 alternative phylogenetic trees representing uncertainty in species placement. All phylogenetic comparative analyses were repeated across these trees, and parameter estimates were summarized as the mean and standard deviation across trees.

### Predicting fresh trait values

To evaluate whether fresh-tissue trait values can be predicted from herbarium-based measurements, we fitted Random Forest (RF) and Gradient Boosting (GB) regression models independently for each trait using species-level mean data, implemented in the R packages randomForest (Liaw and Wiener, 2002) and gbm (Ridgeway, 2013), respectively. For each trait, the dataset was randomly partitioned into training (80%) and testing (20%) sets using a fixed seed to ensure reproducibility. Random Forest models were fitted with 1000 trees and the number of variables considered at each split set to the square root of the total number of predictors. GBM models were fitted with 3000 trees, a maximum tree depth of 3, a learning rate of 0.01, and a subsampling fraction of 0.7. Predictive performance was evaluated on the test data using root mean squared error (RMSE), normalized RMSE (NRMSE; RMSE divided by the trait range), and the coefficient of determination (R^2^).

To assess whether additional information improved prediction performance, we incorporated phylogenetic information and climatic variables in increasingly complex model formulations: (i) herbarium trait measurements only; (ii) herbarium traits combined with phylogenetic information; (iii) and herbarium traits combined with both phylogenetic and climatic variables. Phylogenetic information was incorporated using phylogenetic eigenvectors derived from phylogenetic trees through principal coordinate analysis of cophenetic distances on the dated tree, using the R package ape (Paradis and Schliep, 2019). Axes that together explained up to 80% of the variance (limited to ten axes) were retained. To account for uncertainty associated with random species insertion in the phylogeny, analyses were repeated across all alternative trees, and the mean and standard deviation of model performance metrics were calculated across trees (details of species insertion are provided earlier in the Methods). Climatic variables were included as species-level means of mean annual temperature, temperature seasonality, annual precipitation, and precipitation seasonality at specimen collection sites. Geographic coordinates of each specimen were used to extract climatic data from WorldClim 2.0 (Fick and Hijmans, 2017) using the R packages raster and geodata (Hijmans et al., 2015, 2024).

## RESULTS

### Herborization effect and water content

The effects of herborization varied among plant organs (Fig. 1). Fruit traits tended to exhibit positive effects, whereas most leaf and floral size traits had negative values. Seed traits had mean effects close to zero. Although reductions were observed in leaf size metrics, shape-related traits, such as length-to-width ratio and circularity index, as well as leaf mass per area, generally increased after herborization. Variation in herborization effects among species was comparatively low for leaf traits and more pronounced for fruit traits, with floral and seed traits having intermediate levels of variation. When the herborization effect for fruits was decomposed into pressing and drying components (Fig. S3), fruit length and width increased during pressing but subsequently decreased during drying, resulting in the net positive effect observed.

**Figure 1.**
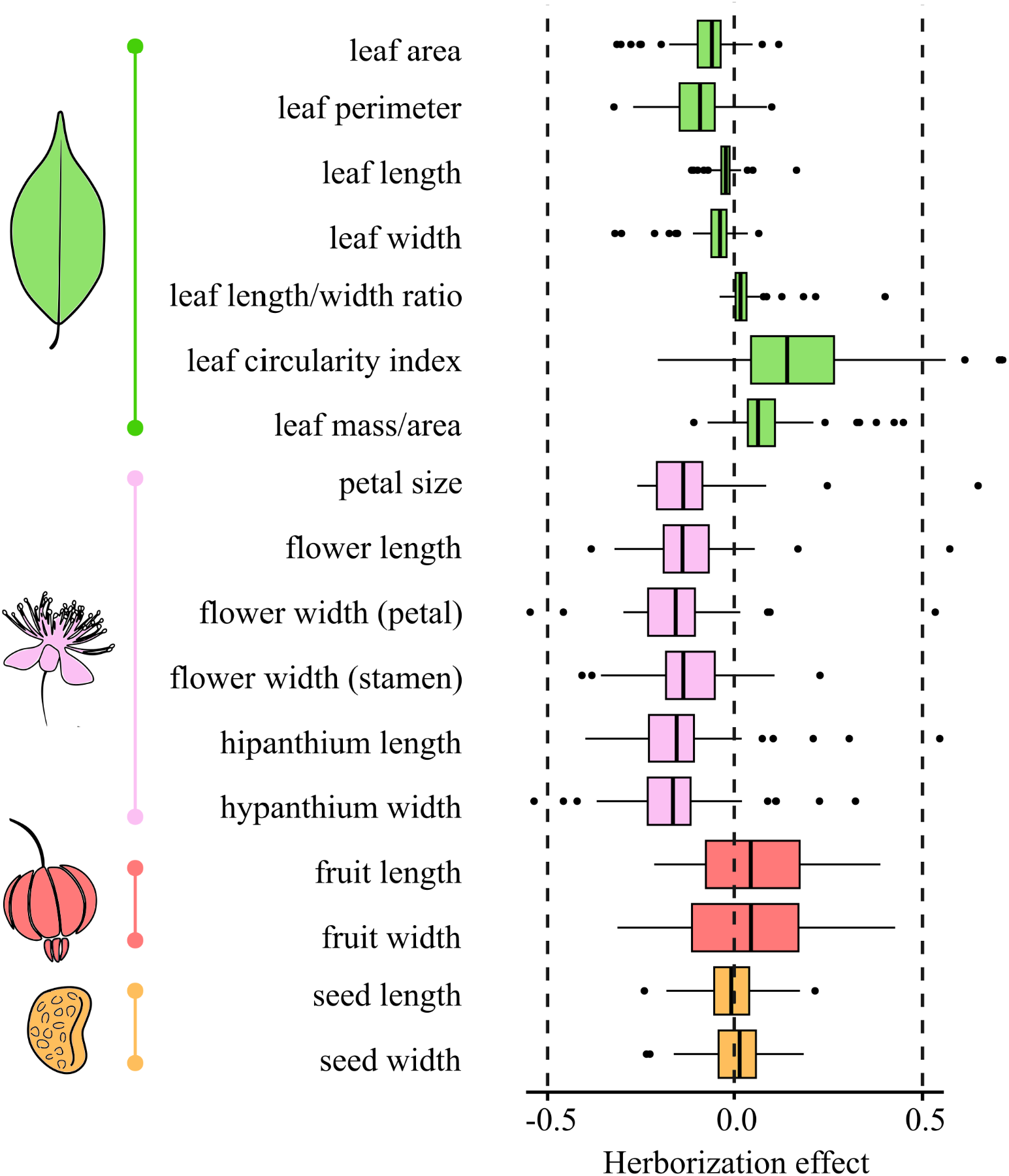
Herborization effect across leaf, flower, fruit, and seed traits. Positive values indicate an increase in trait measurements, whereas negative values indicate a decrease after herborization.

Water content explained a variable proportion of the herborization effect across plant organs (Fig. 2; Table 1). Significant associations between water content and the herborization effect were generally negative, indicating greater shrinkage in structures with higher water content. The only exception was leaf mass per area, which showed a positive relationship, indicating that leaves with higher water content tended to exhibit greater increases in leaf mass per area after herborization. For fruits, no significant relationships were detected (Fig. 2; Table 1); however, trends were positive, suggesting that the herborization effect tends to result in greater expansion in fruits with higher water content. The herborization effect on flower hypanthium dimensions and leaf shape-related traits showed weak or non-significant relationships with water content (Fig. 2; Table 1).

**Table 1.** Summary statistics for PGLS models across the 100 alternative phylogenetic trees (mean and sd) testing the relationship between trait herborization effect and water content for each plant organ.

| Trait | Slope (mean) | Slope (sd) | p value (mean) | R2 (mean) | R2 (sd) |
| --- | --- | --- | --- | --- | --- |
| Flower width (petal) | -0,24384026 | 0,01449239 | 0,01207981 | 0,19691979 | 0,05530127 |
| Flower width (stamen) | -0,39214905 | 0,02181113 | 0,00669024 | 0,21556878 | 0,04568122 |
| Hypanthium width | 0,15725916 | 0,01339637 | 0,26586882 | 0,05434366 | 0,04193678 |
| Hypanthium length | -0,29219894 | 0,00660988 | 0,05363341 | 0,13210540 | 0,04222062 |
| Flower length | -0,79537896 | 0,00046846 | 0,00004075 | 0,40418720 | 0,00109989 |
| Petal size | -0,52045398 | 0,00281888 | 0,00000044 | 0,56758677 | 0,00653224 |
| Fruit width | 0,02853937 | 0,05136223 | 0,85774145 | 0,32429006 | 0,02518732 |
| Fruit length | 0,18628523 | 0,01515661 | 0,41646934 | 0,17719487 | 0,01664554 |
| Leaf area | -0,18762177 | 0,00526459 | 0,00137893 | 0,19731026 | 0,02569370 |
| Leaf circularity index | 0,18048391 | 0,00143589 | 0,29584891 | 0,00986783 | 0,00583313 |
| Leaf length | -0,04137087 | 0,00195487 | 0,13170018 | 0,09599497 | 0,02422306 |
| Leaf length/width | 0,11138431 | 0,02338754 | 0,15559992 | 0,07683805 | 0,08232525 |
| Leaf mass per area | 0,24305401 | 0,00645281 | 0,00125285 | 0,21545671 | 0,01657239 |
| Leaf perimeter | -0,15578900 | 0,00420026 | 0,03004510 | 0,05195782 | 0,02342023 |
| Leaf width | -0,12282659 | 0,00276052 | 0,00853460 | 0,05840962 | 0,01347688 |
| Seed width | -0,09666408 | 0,00729293 | 0,26367383 | 0,19616219 | 0,02061867 |
| Seed length | -0,18806221 | 0,00391914 | 0,01303492 | 0,21106978 | 0,01357359 |

**Figure 2.**
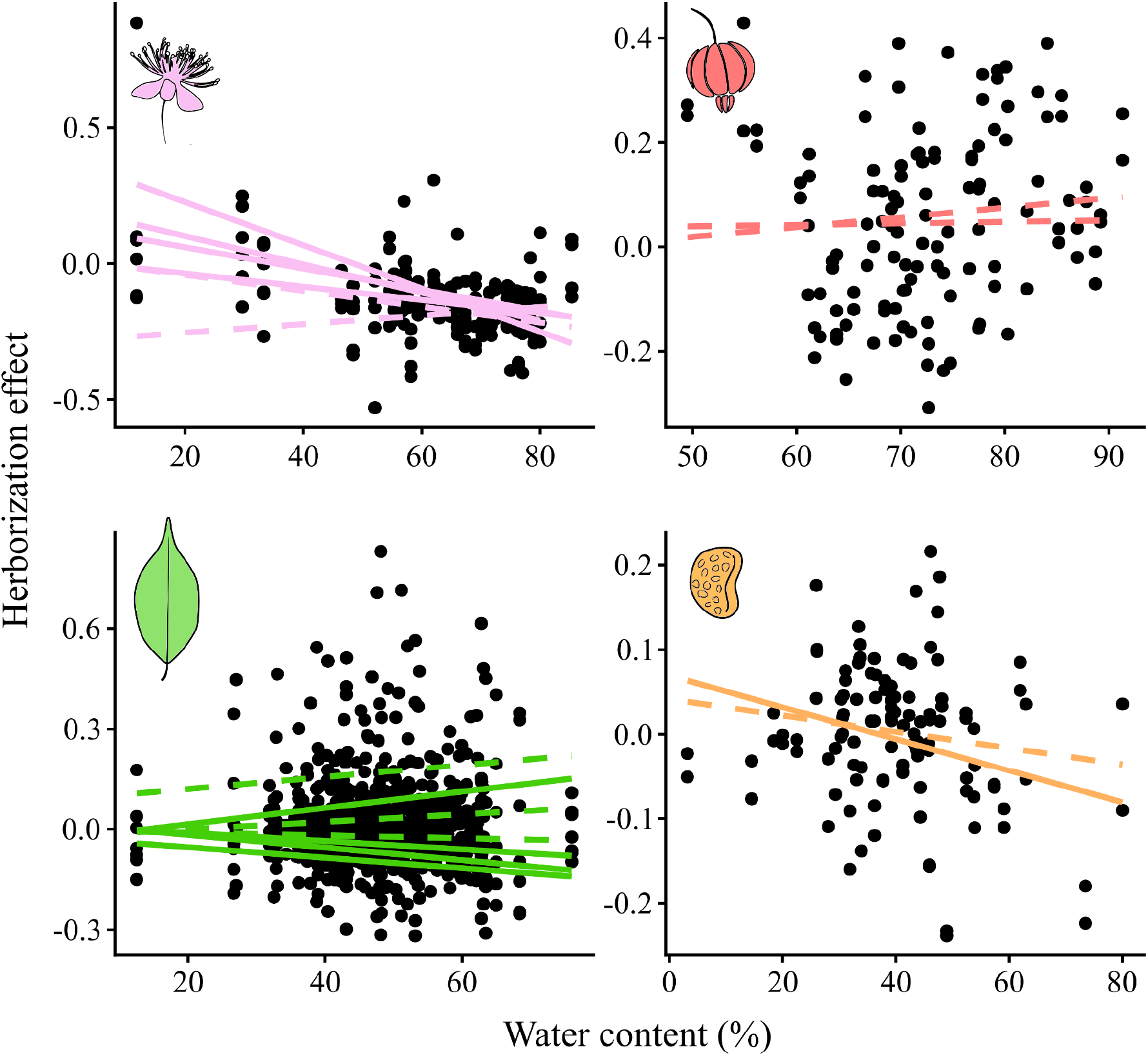
Effect of water content on the herborization effect for flower, fruit, leaf, and seed traits. Each point represents the mean trait value for a species. Lines indicate slopes estimated by PGLS models for each trait; dashed lines represent non-significant relationships.

### Predicting fresh tissue trait values

Models based on herbarium measurements successfully predicted fresh trait values across most plant structures (Fig. 3, Table S2). For most traits, the best-performing model (lowest NRMSE and higher R^2^) was the one using only dried herbarium measurements.

**Figure 3.**
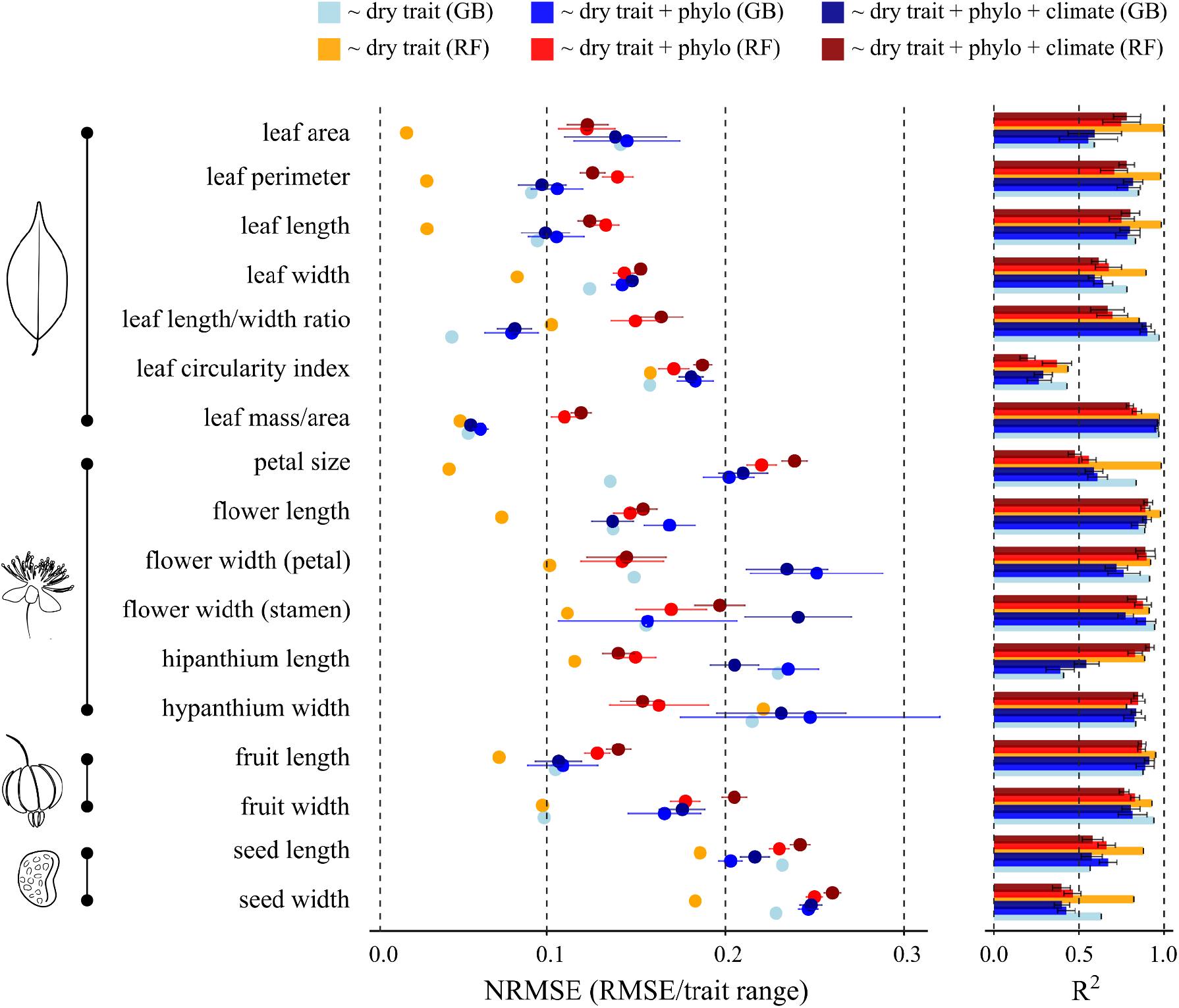
Comparison of model performance for Gradient Boosting (GB) and Random Forest (RF) regression models predicting fresh-tissue trait values from herbarium-based measurements and additional predictors. Panels show normalized root mean squared error (NRMSE; left) and the coefficient of determination (R^2^; right) for each trait. Confidence intervals represent the standard deviation across analyses using 100 alternative phylogenetic trees. Models differ in predictor sets: herbarium-based measurements only (~ dry trait), herbarium traits with phylogenetic information (~ dry trait + phylo), and herbarium traits with both phylogenetic information and climatic variables (~ dry trait + phylo + climate). Trait categories (leaf, flower, fruit, and seed) are indicated by icons on the left.

Random Forest (RF) performed better in general, while Gradient Boosting (GB) had better predictive performance for leaf circularity index, hypanthium width, and seed length and width. In a few cases, models including additional predictors performed best: the phylogeny-informed model for leaf width and leaf circularity index, and the model including both phylogenetic and climatic predictors for hypanthium and fruit width (Fig. 3, Table S2).

Leaf traits showed high predictive performance in the selected models (Fig. 3, Table S2). Leaf size metrics and leaf mass per area were accurately predicted (R^2^ = 0.89–0.99; NRMSE = 0.02–0.08), whereas leaf shape metrics showed lower predictive performance (R^2^ = 0.43–0.87; NRMSE = 0.12–0.16). Floral traits were also accurately predicted (R^2^ = 0.95–0.98; NRMSE = 0.05–0.11), with the exception of hypanthium measurements, which showed comparatively lower predictive performance (R^2^ = 0.85–0.91; NRMSE = 0.12–0.15). Fruit traits were accurately predicted from herbarium measurements (R^2^ = 0.92–0.95; NRMSE = 0.07–0.11). In contrast, seed traits showed lower predictive performance (R^2^ = 0.82–0.88; NRMSE = 0.18–0.19).

Overall, NRMSE and R^2^ values indicate that dried trait measurements are strong predictors of fresh values across most traits. Phylogenetic information and climatic variables provided little additional reduction in prediction error (Fig. 3, Table S2), indicating that dried trait values alone are generally the best predictors of fresh tissue trait values.

## DISCUSSION

This study evaluated how herborization affects morphological trait measurements and whether fresh trait values can be reliably inferred from herbarium specimens across multiple organs in Neotropical Myrtaceae, with potential applicability to other fleshy-fruited woody taxa. We hypothesized that (H1) morphological change during herborization would increase with organ water content and that (H2) fresh trait values could be accurately predicted from herbarium-based measurements. Our findings provide partial support for the first hypothesis, as water content was strongly associated with herborization effects primarily in floral traits but showed weaker or non-significant relationships in leaves, fruits, and seeds. In contrast, the second hypothesis was strongly supported: measurements obtained from herbarium specimens accurately predicted fresh trait values across most traits. Together, these results indicate that although herborization can alter some morphological measurements, particularly in water-rich tissues, herbarium specimens retain substantial quantitative information that can be used to estimate fresh trait values.

### Herborization effect and water content

Herborization effects varied among plant structures. The reductions observed in leaf dimensions are consistent with previous studies showing that drying commonly causes leaf shrinkage (Blonder et al., 2012; Tomaszewski and Górzkowska, 2016). The increase in leaf mass per area after drying also aligns with earlier findings that specific leaf area (the inverse of leaf mass per area) decreases following herborization due to reductions in leaf area during desiccation (Queenborough and Porras, 2014; Perez et al., 2020). Because both metrics are calculated using leaf dry mass (Pérez-Harguindeguy et al., 2016), a decrease in leaf area necessarily leads to higher leaf mass per area and lower specific leaf area. The magnitude of shrinkage observed here is comparable to that reported for woody and evergreen taxa (approx. 10%; Blonder et al., 2012), which is particularly relevant given the predominance of woody evergreen species in Myrtaceae. Water content was generally negatively associated with the magnitude of herborization effects in size-related leaf traits, but these relationships were weak, suggesting that structural properties such as lignin content (Juneau and Tarasoff, 2012) may play a more important role in determining leaf shrinkage.

Despite the general shrinkage of leaf dimensions, shape-related traits such as length-to-width ratio and circularity index increased after herborization. This pattern suggests that desiccation affects leaf axes unevenly, resulting in differential shrinkage rather than uniform contraction. Therefore, although herbarium specimens provide reliable estimates of absolute leaf size, caution is needed when interpreting shape-based traits that are sensitive to proportional changes. Tomaszewski and Górzkowska (2016) similarly reported that leaf shape is often not fully preserved after drying. In most species they examined, leaves became narrower, consistent with our observed increase in length-to-width ratio. However, the magnitude of these alterations appeared highly species-specific, which the authors attributed to anatomical differences, particularly in secondary venation patterns and lignin content.

Notably, that study was based on relatively limited taxonomic sampling (22 species), highlighting the need for broader comparative assessments across diverse lineages.

Extending this framework beyond leaves, our study shows that floral traits exhibit similar but often stronger size reductions. Flowers tend to be less lignified and are typically short-lived organs not adapted for structural persistence, which likely explains their greater sensitivity to desiccation (Roddy et al., 2023; Ievinsh, 2023). Consistent with this, floral traits showed the strongest and most consistent relationships with water content, with species exhibiting higher floral water content experiencing greater shrinkage during drying. This pattern supports the expectation that tissues with a higher proportion of hydrated cells are particularly sensitive to desiccation.

Seed traits, in contrast, exhibited relatively limited dimensional changes. This may reflect the fact that dehydration is an intrinsic component of seed maturation, meaning their tissues are developmentally adapted to water loss (Bewley et al., 2012). Although most Myrtaceae seeds are recalcitrant and retain relatively high water content compared with seeds in many other plant clades (Wyse and Dickie, 2017), the magnitude of shrinkage observed here was small, suggesting that seed measurements from herbarium specimens may still be broadly reliable. Consistent with this interpretation, seed traits showed intermediate relationships with water content, indicating that hydration contributes to variation in shrinkage but is not its primary driver.

Fruit traits showed no clear relationship between water content and herborization effects. The absence of a clear water-content signal in fruits may reflect the strong mechanical effects associated with specimen preparation. This indicates that dimensional changes in fruits are largely driven by mechanical flattening prior to desiccation rather than by water loss alone. Fruits in the tribe Myrteae are predominantly fleshy berries with high water content, often ranging from 71 to 94% of pulp mass (Galetti et al., 2011). Pressing these three-dimensional structures during specimen preparation can therefore substantially distort their morphology, particularly in mature fruits, demonstrating that herborization effects may reflect multiple processing steps rather than drying alone.

To our knowledge, this study is the first to explicitly disentangle and quantify the separate contributions of pressing and drying to herborization-related measurement bias. While this distinction may be less critical for relatively flat structures such as leaves (e.g., Juneau and Tarasoff, 2012; Blonder et al., 2012; Queenborough and Porras, 2014; Tomaszewski and Górzkowska, 2016; Perez et al., 2020), it is particularly important for fleshy and three-dimensional organs. These results therefore indicate that fruit traits derived from herbarium specimens require particular caution, especially in clades characterized by fleshy fruits with high water content.

### Predicting fresh tissue trait values

Fresh trait values were reliably predicted from herbarium-based measurements across most plant structures. In most cases, models using only dried trait measurements performed best, indicating that dimensional changes caused by herborization are sufficiently consistent to allow accurate reconstruction of fresh values without additional predictors.

Predictive performance was highest for leaf size traits, reinforcing previous findings that herbarium specimens provide reliable estimates of leaf dimensions (Blonder et al., 2012). In contrast, leaf shape metrics were less accurately predicted, likely reflecting uneven deformation during drying that alters proportional relationships among leaf axes (Tomaszewski and Górzkowska, 2016). Floral traits also showed strong predictive performance overall, although some structures such as the hypanthium were slightly more sensitive to distortion during specimen preparation. Predictions for fruits and seeds were generally robust but somewhat less accurate, probably reflecting the three-dimensional structure of these organs and the mechanical effects associated with pressing these tissues.

Additional predictors contributed little to model performance. Phylogenetic information improved predictions for only a few traits, and climatic variables rarely reduced prediction error. Together, these results indicate that dried trait measurements alone usually capture most of the information needed to estimate fresh values. This pattern aligns with the conclusions of Blonder et al. (2012), who found that evolutionary history and climate played minor roles when predicting leaf shrinkage.

### Concluding remarks

Our findings reinforce the potential of herbarium collections as large-scale sources of biological trait data. The ability to reliably infer fresh morphological traits from preserved specimens supports emerging collectomics approaches (Bucher et al., 2025), which aim to extract and integrate biological information from natural history collections using automated and computational tools. Because collecting fresh trait data in the field often requires substantial logistical effort and financial investment, particularly in remote or species-rich regions, herbarium specimens provide a powerful complementary resource for expanding trait datasets across taxa, space, and time (Davis, 2023). Our results therefore contribute to the growing role of collectomics in leveraging biological collections for large-scale ecological and evolutionary research (Sigwart et al., 2025).

## ACKNOWLEDGMENTS

The authors thanks Bispo MRB, Campelo V, Carvalho MS, Castro CO, Cunha HF, Faria JE, Haluch CA, Marujo B, Meyer FS, Moreira VP, Rosário AS, Souza MC, Souza PJS, and Vendramini JV for their invaluable assistance during field expeditions. We are also grateful to Vieira BM for help with figure design. We thank the staff and curators of the herbaria ALCB, CEPEC, IAN, MBM, MG, RB, SPF, UB, and UFRN for logistical support. This study was financed in part by the Coordenação de Aperfeiçoamento de Pessoal de Nível Superior - Brasil (CAPES) - Finance Code 001 and by the Instituto Serrapilheira (grant number Serra-R-2111-39858). We also acknowledge the Sociedade Botânica do Brasil (SBB) for financial support through the Scientia Amabilis 2024 grant.

## AUTHOR CONTRIBUTIONS

**Yacov Kilsztajn** Conceptualization (equal); formal analysis (lead); investigation (lead); methodology (lead); visualization (lead); writing – original draft preparation (lead); writing – review & editing (equal). **Lázaro Henrique Soares de Moraes Conceição:** Investigation (supporting); writing – review & editing (equal). **Thais Vasconcelos:** Conceptualization (equal); investigation (supporting); writing – review & editing (equal). **Carolyn Elinore Barnes Proença:** Investigation (supporting); writing – review & editing (equal). **Vanessa Staggemeier:** Conceptualization (equal); investigation (supporting); funding acquisition (lead); resources (lead); supervision (lead); writing – review & editing (equal).

## SUPPORTING INFORMATION AND DATA AVAILABILITY

Supporting Information and the code used to perform the analyses are available at https://github.com/ykilsztajn/fresh_dry_myrtaceae. All raw data will be made available in the same repository upon acceptance for publication.

**Figure S1.** Geographic distribution of sampled specimens across Brazil. Sampling localities within a 0.3° radius were aggregated for visualization purposes.

**Figure S2.** Details of flower trait measurements: (a) petal size; (b) flower length; (c) flower width (petal-to-petal); (d) flower width (stamen-to-stamen); (e) hypanthium length; (f) hypanthium width.

**Figure S3.** Separate effects of fruit pressing and drying on length and width measurements.

**Table S1.** Sampled specimens with corresponding herbarium vouchers and information on whether each species is included in the most recently published phylogeny for Neotropical Myrtaceae (NMWG et al., 2024).

**Table S2.** Comparison of model performance for Gradient Boosting (GB) and Random Forest (RF) regression models predicting fresh-tissue trait values from herbarium-based measurements. Summary performance metrics are reported for each trait across all models and across the 100 alternative phylogenetic trees (mean and sd). Models differed in predictor sets: herbarium-based measurements only (~ dry trait), herbarium traits plus phylogenetic information (~ dry trait + phylo), and herbarium traits plus both phylogenetic information and climatic variables (~ dry trait + phylo + climate).

